# Acute Hyperalgesia and Delayed Dry Eye After Corneal Abrasion Injury

**DOI:** 10.1101/242685

**Authors:** Deborah M. Hegarty, Sam M. Hermes, Michael M. Morgan, Sue A. Aicher

## Abstract

Corneal nerves mediate pain from the ocular surface, lacrimation, and blinking, all of which protect corneal surface homeostasis and help preserve vision. Corneal nerve density correlates with neuropathic pain states and is used as an assessment of small fiber neuropathies. Because pain, lacrimation and blinking are rarely assessed at the same time, it is not known if their regulatory mechanisms have similar temporal dynamics after acute corneal injury. We examined changes in corneal nerve density, evoked and spontaneous pain, and ocular homeostasis in Sprague-Dawley male rats after a superficial epithelial injury with heptanol that acutely abolished nerve endings within the central cornea. Despite a profound loss of epithelial nerve endings, pain was transiently enhanced after abrasion injury, while basal tear production was normal. We found no relationship between epithelial nerve density and pain or homeostatic responses. Axotomy following corneal abrasion increased expression of both ATF3 (a nerve injury marker) and CGRP (a nociceptive peptide) in trigeminal ganglia 24 hours after injury. These molecular changes were absent on the contralateral side, despite reductions in corneal epithelial nerve density in the uninjured eye. ATF3 and CGRP levels in trigeminal ganglion were normal at one week post-injury when pain responses were normal. In contrast, CGRP was upregulated in peripheral corneal endings one week after injury, when dry eye symptoms emerged. Our results demonstrate dynamic trafficking of CGRP within trigeminal sensory nerves, with elevations in the ganglion correlated with pain behaviors and elevations in peripheral endings correlated with dry eye symptoms.

## 1. Introduction

The cornea contains the highest density of sensory nerves of any tissue in the body [3; 39; 42]. Corneal sensory nerves are predominantly nociceptive and even low threshold stimulation evokes a sensation of pain [3]. Corneal nerves are also vital for homeostatic reflexes, such as blinking and lacrimation, and epithelial health [6]. Loss of innervation causes a loss of sensation [47] and the development of spontaneous corneal abrasions [55]. Damage to corneal nerves can be caused by diabetes [29], pharmacotherapeutics such as paclitaxel [14], vision correction surgery [37], or accidental injury [20] that causes damage to the ocular surface. Studies examining injury-induced changes in corneal nerves find conflicting and paradoxical results when trying to correlate altered nerve structure to ocular pain. In some cases ocular sensation is unchanged despite obvious corneal nerve damage, and in other cases persistent pain may occur [3; 9]. The presence of ocular pain may be caused by neuropathic changes in corneal nerves. The present study tested this hypothesis by inducing a unilateral corneal abrasion injury in rats to look at the relationship between changes in corneal nerve density and morphology, neurochemistry, and ocular sensation. We examined temporal changes in ocular homeostatic responses, including blinking and tear production. We also examined molecular changes in corneal sensory cell bodies and peripheral endings, using activating transcription factor 3 (ATF3) as a marker of nerve injury [7] and calcitonin gene-related peptide (CGRP) as a molecule associated with changes in nociceptive processing [8] and epithelial health [51].

We found distinct temporal dynamics for changes in nociception versus homeostatic ocular responses. Pain increased acutely after superficial corneal epithelial injury, but pain responses returned to normal levels long before nerve density returned to normal; in contrast, tear production was normal acutely but was impaired one week after corneal damage when re-innervation of the corneal epithelium is progressing and pain responses are normal. Together these findings suggest divergent mechanisms underlying ocular pain and homeostatic functions, even though sensory nerves and some molecular transducers such as TRPM8 are necessary for both cold pain [4] and homeostatic responses [48]. The trigeminal sensory neurons that innervate the corneal surface are molecularly complex and may differentially traffic materials to central and peripheral sites. These neurons are also integrated into distinct reflex pathways that mediate pain responses, blinking and lacrimation.The present studies indicate that trafficking of the neuropeptide calcitonin gene related peptide (CGRP) to different regions of the sensory nerves innervating the cornea is correlated with distinct functional changes in nociceptive and homeostatic responses.

## 2. Methods

Methods were approved by Institutional Animal Care and Use Committee (IACUC) at OHSU. Experiments were performed on male Sprague-Dawley rats (250–350 g). Rats used for wheel running studies were bred at Washington State University and were housed individually. Rats studied at OHSU (Charles River Laboratories, Wilmington, MA) were housed in pairs on a 12/12 light/dark cycle and had access to food and water at all times. A total of 52 rats were used for all of the experiments, specific groups sizes are indicated in the Results section for each study. In the course of measuring behavioral responses, corneal nerve density, and homeostatic responses, some rats at OHSU were assessed for more than one metric when possible.

### 2.1. Heptanol abrasions & tracer applications

Rats were anesthetized with isoflurane for unilateral corneal abrasion. Heptanol (10 μl) was pipetted into a small metal ring (6-gauge (4.4 mm internal diameter (ID)) stainless steel tubing, custom, HTX-06R, Small Parts Inc., Miramar, FL) secured to the cornea with petroleum jelly and left for 90 seconds [19; 24]. After 90 seconds, the heptanol was wicked away with cotton swabs, a cotton-tipped applicator was used to remove the damaged epithelium from the central cornea and the corneal surface was profusely rinsed with saline. Rats were returned to their home cage and monitored until they were awake from anesthesia.

In a subset of animals, the tract tracer FluoroGold (FG, Fluorochrome, Denver, CO) was applied to the central cornea in the 6-gauge metal ring immediately following the heptanol abrasion to allow labeling of nerves that were severed during the initial abrasion. One week after abrasion, 1% Cholera toxin B (CTb; List Laboratories, Campbell, CA) was applied to the cornea using a similar method, but with a smaller metal ring (7-gauge (3.8 mm ID), HTX-07R, Small Parts, Inc.). For both tract tracer applications, the dye was left on the cornea for 30 minutes [22]. Animals were perfused 1 week after CTb application as described below and trigeminal ganglia (TG) were examined for the presence of single (CTb)- or dual (CTb and FG)-labeled neurons (see section 2.5).

### 2.2. Evoked and spontaneous pain measures: Menthol-evoked eye wipes and spontaneous wheel running

To assess evoked pain responses, menthol (10 μl of 50 mM menthol in 10% ethanol/10% Tween-80 in saline) was pipetted directly onto the corneal surface in awake, lightly restrained rats. Two observers counted the number of ipsilateral eye wipes for 3 minutes after menthol stimulation [23]. Rats were habituated to handling, the testing apparatus and testing room for 40 minutes per day for 2 days prior to nociceptive testing. Each rat received a single corneal menthol application to avoid sensitization; thus, separate groups were used for 24 hour and 1 week assessments. Individual rats were tested either on the abraded or contralateral eye, but not both.

Home cage wheel running was used to assess spontaneous pain as described previously [31; 32]. Rats were given access to a running wheel in the home cage for 23 hours a day for 7 days prior to corneal abrasion. The number of wheel revolutions measured on the seventh day was used as the baseline value. Rats were randomly assigned to control or abrasion groups. Only rats with a baseline wheel running value of at least 400 revolutions on the baseline day were tested. Testing began approximately 1 hour after induction of a unilateral heptanol abrasion of the cornea as described above. Control rats were briefly anesthetized but not abraded. Rats were returned to the home cage and wheel running was assessed 23 hours a day for one week. No one entered the test room except during the one hour when running wheel data was not assessed. Food, water, and the health of the rat was monitored during this hour.

### 2.3. Spontaneous Blinks and Winks

The number of blinks (simultaneous eye closures of both eyes) and winks (unilateral eye closures) were counted prior to (Baseline) and 1 week after corneal abrasion. Rats were placed in a Plexiglas chamber and allowed to acclimate for 15 minutes. Blinks and winks were counted by an observer for 5 minutes. Rats were habituated to handling, the testing apparatus, and testing room for 60 minutes per day for 2 days prior to eye closure assessment.

### 2.4. Phenol thread test

Tear production was assessed using the phenol thread test (Zone-Quick, Oasis Medical, Glendora, CA) as described previously [1]. This test is equivalent to the Schirmer’s test [53]. A timed wicking (15 seconds/eye) was used to measure tear production in rats briefly anesthetized with isoflurane (5% in oxygen). At the end of 15 seconds, the length of thread that turned red was measured to the nearest millimeter (mm). Each animal was tested prior to abrasion (Baseline) and at a time point post-abrasion (24 hours; 1 week). Phenol thread test was done at least 24 hours prior to nociceptive testing because we have observed that even brief anesthesia interferes with nociceptive testing on the same day.

### 2.5. Immunohistochemistry

Rats were overdosed with sodium pentobarbital (150 mg/kg) and perfused transcardially through the ascending aorta with 10 ml of heparinized saline (1000 units/ml) followed by 600 ml of 4% paraformaldehyde in 0.1 M phosphate buffer, pH 7.4 (PB) [23; 26]. The eyes were enucleated immediately after perfusion and placed in PB. Using a 5 mm corneal trephine (Ambler Surgical, Exton, PA), the central corneas were removed and placed in fresh PB at 4 °C until immunoprocessing. The trigeminal ganglia were dissected, placed in fixative for 30 minutes, then stored in 30% sucrose in PB for at least 24 hours. Trigeminal ganglia were then cryosectioned at 20 μm on a Leica CM1950 cryostat (Leica Microsystems, Inc. Buffalo Grove, IL) and mounted directly onto room temperature Superfrost® Plus slides (Fisher Scientific, Pittsburgh, PA). Sections were dried and stored at -20 °C until immunoprocessing.

Whole mount corneas were processed free-floating while trigeminal ganglia were processed on slides as previously described [1; 22; 23]. Tissue was incubated in 1% sodium borohydride solution for 30 minutes prior to primary antibody incubation. Corneas were incubated for 3 nights and trigeminal ganglia sections were incubated for 2 nights at 4 °C in primary antibody cocktails in 0.25% Triton X-100 (Sigma-Aldrich, St. Louis, MO) and 0.1% Bovine Serum Albumin (BSA, Sigma) in 0.1 M Tris-buffered saline pH 7.6 (TS) (Table 1). Corneas and trigeminal ganglia sections were rinsed and incubated in fluorescent secondary antibody cocktails made in 0.1% BSA in TS (Table 1) light-protected for 2 hours at room temperature. Corneas were rinsed, blotted dry and placed into individual wells of an 18 well μ-slide (ibidi USA, Inc., Fitchburg, WI) and coverslipped with CFM-1 (Electron Microscopy Sciences, Hatfield, PA) which maintains the natural curvature of the cornea for imaging. Trigeminal ganglia sections were additionally stained with NeuroTrace Fluorescent Nissl stain 530/615 (1:200, ThermoFisher Scientific, Waltham, MA) for 20 minutes before being rinsed, dried and coverslipped with Prolong Gold™ Antifade reagent (Life Technologies).

**Table 1.**
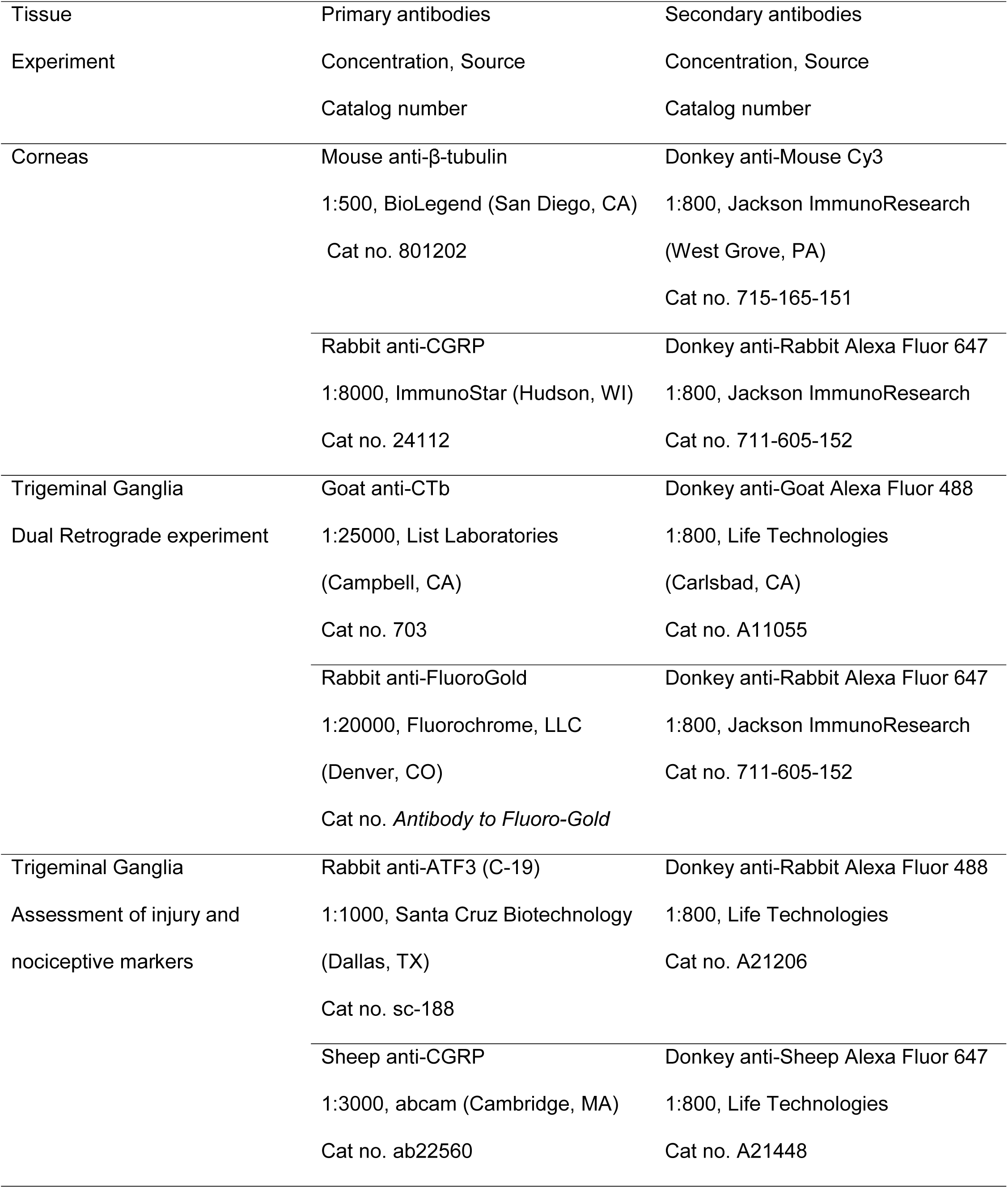
List of primary and secondary antibodies used for each immunohistochemical experiment.

| Tissue | Primary antibodies | Secondary antibodies |
| --- | --- | --- |
| Experiment | Concentration, Source | Concentration, Source |
|  | Catalog number | Catalog number |
| Corneas | Mouse anti-β-tubulin | Donkey anti-Mouse Cy3 |
|  | 1:500, BioLegend (San Diego, CA) | 1:800, Jackson ImmunoResearch |
|  | Cat no. 801202 | (West Grove, PA) |
|  |  | Cat no. 715-165-151 |
|  | Rabbit anti-CGRP | Donkey anti-Rabbit Alexa Fluor 647 |
|  | 1:8000, ImmunoStar (Hudson, WI) | 1:800, Jackson ImmunoResearch |
|  | Cat no. 24112 | Cat no. 711-605-152 |
| Trigeminal Ganglia | Goat anti-CTb | Donkey anti-Goat Alexa Fluor 488 |
| Dual Retrograde experiment | 1:25000, List Laboratories | 1:800, Life Technologies |
|  | (Campbell, CA) | (Carlsbad, CA) |
|  | Cat no. 703 | Cat no. A11055 |
|  | Rabbit anti-FluoroGold | Donkey anti-Rabbit Alexa Fluor 647 |
|  | 1:20000, Fluorochrome, LLC | 1:800, Jackson ImmunoResearch |
|  | (Denver, CO) | Cat no. 711-605-152 |
|  | Cat no. *Antibody to Fluoro-Gold* |  |
| Trigeminal Ganglia | Rabbit anti-ATF3 (C-19) | Donkey anti-Rabbit Alexa Fluor 488 |
| Assessment of injury and nociceptive markers | 1:1000, Santa Cruz Biotechnology | 1:800, Life Technologies |
|  | (Dallas, TX) | Cat no. A21206 |
|  | Cat no. sc-188 |  |
|  | Sheep anti-CGRP | Donkey anti-Sheep Alexa Fluor 647 |
|  | 1:3000, abcam (Cambridge, MA) | 1:800, Life Technologies |
|  | Cat no. ab22560 | Cat no. A21448 |

### 2.6. Imaging and Image analysis

#### 2.6.1. Corneal nerve density and CGRP content

Antibodies to β-tubulin and CGRP (Table 1) were used to assess corneal nerve density and CGRP content as previously described [23]. Images of β-tubulin- and CGRP-labeled corneal nerves were taken within the central cornea and included the whorl-like vortex as an anchor. Images were captured on a Zeiss LSM510 or LSM780 confocal microscope with a 20x Plan-Apochromat objective (Carl Zeiss MicroImaging, Thornwood, NY) using the single pass, multi-track format at a 2048 × 2048 pixel resolution. Optical sectioning produced Z-stacks of 1 μm optical slices that were bounded by the extent of fluorescent β-tubulin and CGRP immunolabeling throughout the thickness of the corneal epithelium.

Volumetric assessments of cornea nerve density and CGRP content was performed using Imaris 8.0 software (Bitplane USA, Concord, MA) on an offline workstation in the Advanced Light Microscopy Core at OHSU by a treatment blind observer as previously described in detail [23]. Only corneas that did not receive menthol stimulation were assessed. Briefly, we defined the corneal epithelium by the β-tubulin-and CGRP-labeled sub-basal and intraepithelial corneal nerves from the anterior surface of the cornea to the epithelial-stromal border. The corneal epithelium was isolated as a region of interest (ROI) using the Surfaces Segmentation tool and its volume was calculated by the software (μm^3^). The Mask Channel function was then used to isolate β-tubulin and CGRP labeling within the corneal epithelium ROI for further analysis. β-tubulin labeling within the ROI was determined using the Thresholding function and Surfaces Segmentation tool in Imaris, and a volume (μm^3^) was calculated. In order to account for subtle differences in the corneal epithelium volumes (Table 2), the volume of β-tubulin labeling was calculated as a percentage of the corneal epithelium volume for each cornea and expressed as the mean % β-tubulin or the percent of the epithelium that contains β-tubulin-labeled nerves. To measure the CGRP content specifically within the corneal nerves, we established the β-tubulin volume as the ROI and isolated the CGRP labeling using the Mask Channel function and then CGRP volume (μm^3^) was calculated. The volume of CGRP within the volume of the β-tubulin labeling was calculated as a percentage and expressed as the % β-tubulin containing CGRP.

**Table 2.** Corneal epithelium volumes

| Injury Group | Epithelium volume ($\times 10^6 \mu\text{m}^3$) | N |
| --- | --- | --- |
| Uninjured Control | $6.3 \pm 0.8$ | 6 |
| 24 hours Abraded | n/a | 4 |
| 24 hours Contralateral | $8.0 \pm 0.9$ | 3 |
| 1 week Abraded | $5.7 \pm 0.5$ | 4 |
| 1 week Contralateral | $7.1 \pm 1.1$ | 4 |

Data expressed as mean ± SEM. All volumes have an exponent of $\times 10^6$ .

#### 2.6.2. Trigeminal ganglia

In the dual retrograde experiment, trigeminal ganglia sections were assessed for the number of CTb-labeled corneal neurons and whether they contained FG tracer. Cryostat sections (13 – 15 sections per animal, 120 μm apart) throughout the trigeminal ganglia were evaluated using an Olympus BX51 epifluorescence microscope equipped with a DP71 camera and associated software (Olympus America, Center Valley, PA). All trigeminal ganglia sections with CTb labeling were imaged for CTb, NeuroTrace (NT) and FG labeling. Each marker was imaged separately using a 10x UPlanFL N objective and the appropriate filter (FITC, TRITC, Cy5) for the fluorophore or neuronal stain.

In the assessment of injury and nociceptive markers after abrasion, only one trigeminal ganglion section from Uninjured control animals and one from the ipsilateral and/or the contralateral side of 24 hour and 1 week Abraded animals were evaluated for activating transcription factor 3 (ATF3), a nerve injury marker [7], and CGRP immunocytochemistry. In order to avoid counting corneal neurons activated by noxious corneal menthol stimulation [7], only trigeminal ganglion sections from unstimulated sides were used. Cryostat sections (9 – 14 sections per animal, 120 μm apart) throughout each trigeminal ganglion were first evaluated with the Olympus epifluorescence microscope. The trigeminal ganglion section that contained peak ATF3 labeling per ganglion was also imaged for NT and CGRP using the 10x UPlanFL N objective and the appropriate filter (FITC, TRITC, Cy5). Analysis was performed by an observer blind to the experimental conditions.

Trigeminal ganglion neurons were analyzed using the Cell Counter Plug-In in ImageJ (NIH) similar to our previous study [22]. In order for trigeminal ganglion neurons to be included in each analysis, the nucleus had to be visible in the NT channel. In the dual retrograde experiment, every CTb-labeled trigeminal ganglion neuron was marked with a number using the Cell Counter. The CTb numbers were then transferred to the FG image from the same trigeminal ganglion section and each numbered neuron was evaluated for FG expression. The total number CTb-labeled trigeminal ganglion neurons that contained FG was calculated as a percent for each animal and expressed as % CTb-labeled neurons containing FG.

In order to assess the prevalence of injury and nociceptive markers after abrasion, we confined the analysis to the section that contained the peak number of trigeminal ganglion neurons labeled with the injury marker ATF3. Using the ImageJ Cell Counter Plug-In, every NT-stained trigeminal ganglion neuron with a visible nucleus was marked with a number to get the total number of neurons in that section. Trigeminal ganglion neurons that were ATF3-immunoreactive (-ir) or CGRP-ir were evaluated separately using the Cell Counter Plug-In. The number of trigeminal ganglion neurons that contained ATF3 or CGRP was calculated as a percent for each ganglion and grouped by treatment. Data were expressed as mean % trigeminal ganglion neurons (TGNs) containing ATF3 or CGRP.

### 2.7. Statistics

Statistical analyses and graph production were performed using SigmaPlot 12.0 software. Data are expressed as the mean ± SEM unless otherwise noted. A one-way ANOVA across all treatment groups was performed to test for differences among corneal epithelium volumes. A one-way ANOVA was used to compare eye wipe responses, % β-tubulin, % trigeminal ganglion neurons containing ATF3 and % neurons containing CGRP in the Abraded and Contralateral groups within each post-abrasion time point (24 hour; 1 week) to the Uninjured control group. The Holm-Sidak post hoc test was utilized during these analyses when applicable. We used a t-test to compare % β-tubulin values from the 24 hour Contralateral group and the Uninjured group since there was no β-tubulin to analyze in the 24 hour Abraded group. A t-test was also used to compare baseline wheel running revolution values in Uninjured versus Abraded animals. A nonparametric Mann-Whitney Rank Sum test was used to compare % β-tubulin containing CGRP between Uninjured and 24 hour Contralateral groups while a Kruskal-Wallis one-way ANOVA on Ranks with a Dunn’s post-hoc test was used to compare Uninjured to 1 week Abraded and 1 week Contralateral groups. Paired t-tests were used for within-animal comparisons of raw phenol thread measurements and bilateral and unilateral eye closures measured prior to injury (Baseline) and at time points post-abrasion. A paired t-test or non-parametric Wilcoxon Signed Rank Test was used to test for differences between Baseline and Day 1 wheel revolution values for the Uninjured and Abraded treatment groups, respectively. In all cases, a *P* value less than 0.05 was considered significant.

## 3. Results

### 3.1. Corneal abrasion causes short-term increase in evoked and spontaneous pain

To assess evoked nociceptive responses, menthol was applied to the left eye and the number of nocifensive eye wipes was measured. Eye wipe responses to ocular application of menthol were increased at 24 hours after abrasion injury (Fig. 1; one-way ANOVA (*P* = 0.002)) compared to Uninjured animals (Holm-Sidak post hoc, *P* = 0.004) and the unabraded contralateral eye at this time point (Holm-Sidak post hoc, *P* = 0.006). There was no change in eye wipe responses to menthol in the unabraded, contralateral eye as compared to the Uninjured control eye at 24 hours. Responses in both eyes were at baseline levels one week after injury (Fig.1). The presence of hyperalgesia in the abraded eye is consistent with prior studies of increased photophobia using this model of corneal injury [19].

**Figure 1.**
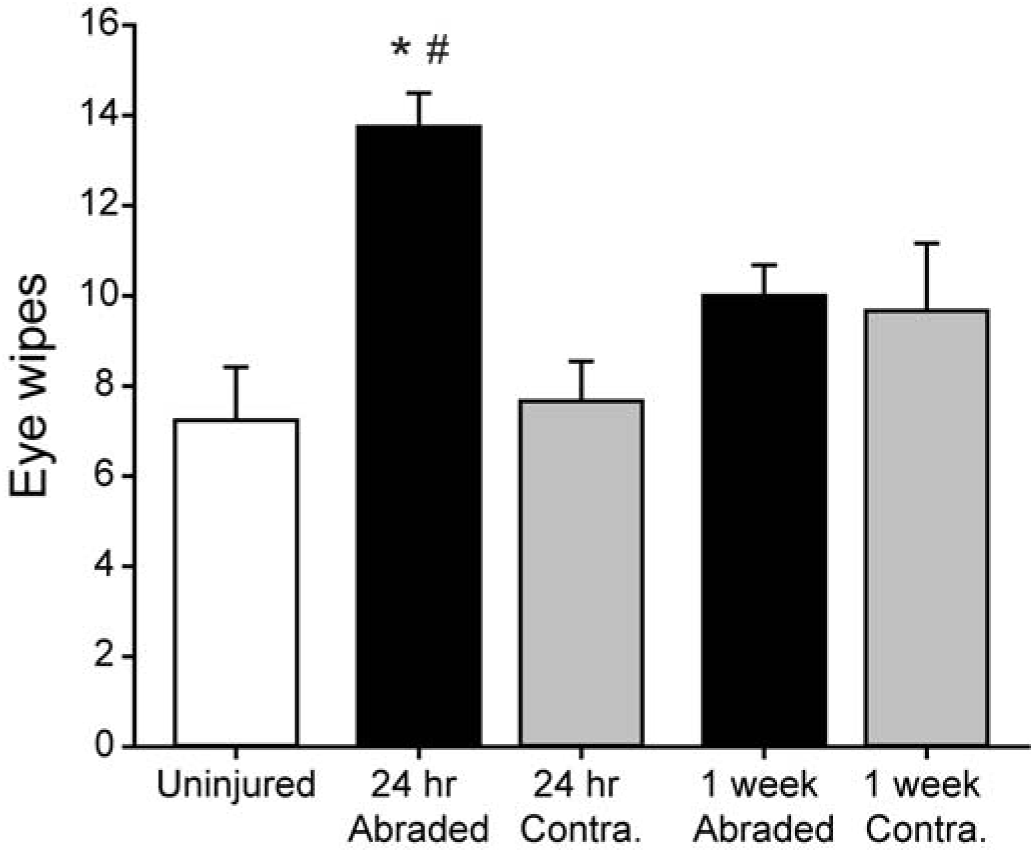
Abraded rats show more eye wipes to ocular application of noxious menthol 24 hours after injury (black bar, n = 4) compared to Uninjured rats (white bar, n = 4) or Abraded rats tested at 24 hours on the contralateral side (24 hr Contra., gray bar, n = 3). At 1 week post abrasion, the eye wipe responses to menthol are normal on both Abraded (black bar, n = 6) and Contralateral eyes (gray bar, n = 6). * *P* < 0.05 versus Uninjured; # *P* <0.05 versus Contralateral side.

There was no difference in baseline wheel running rates between Abraded and Uninjured control animals prior to corneal injury (Abraded baseline: 1418.3 ± 307.7 revolutions; Uninjured baseline: 1169.3 ± 205.9 revolutions; t-test, *P* = 0.306). A significant decrease in wheel revolutions occurred in the 24 hours following corneal abrasion (Abraded 24 hours: 832.8 ± 136.2 revolutions) compared to baseline values (Wilcoxon Signed Rank test, *P* = 0.031), while no change in wheel running was evident in Uninjured control animals (Uninjured 24 hours: 1125.3 ± 394.1 revolutions; Paired t-test, *P* = 0.419). Wheel revolution values returned to baseline levels in Abraded animals by the second day post-abrasion and remained at baseline through one week post-abrasion. These data support the evoked pain data and suggest that spontaneous pain after corneal abrasion peaks 24 hours after injury and resolves by 1 week after corneal injury in Abraded animals.

### 3.2. Central corneal nerves are absent 24 hours after abrasion and partially recover by 1 week

The extent of nerve injury was assessed at times when nociceptive responses were measured in our abraded rats. Using quantitative assessment of β-tubulin immunocytochemistry, we found that heptanol abrasion removed epithelial nerves from the central cornea at 24 hours after injury (Fig. 2). Some corneal innervation returned at 1 week after injury, but nerve density was reduced and did not have the classical central whorl seen in intact corneas (Fig. 2). These findings are intriguing and demonstrate that hyperalgesia can be evoked by ocular application of menthol, even in the absence of epithelial nerve endings in the central cornea. Also, nociceptive responses return to normal at a time when innervation is not fully restored.

**Figure 2.**
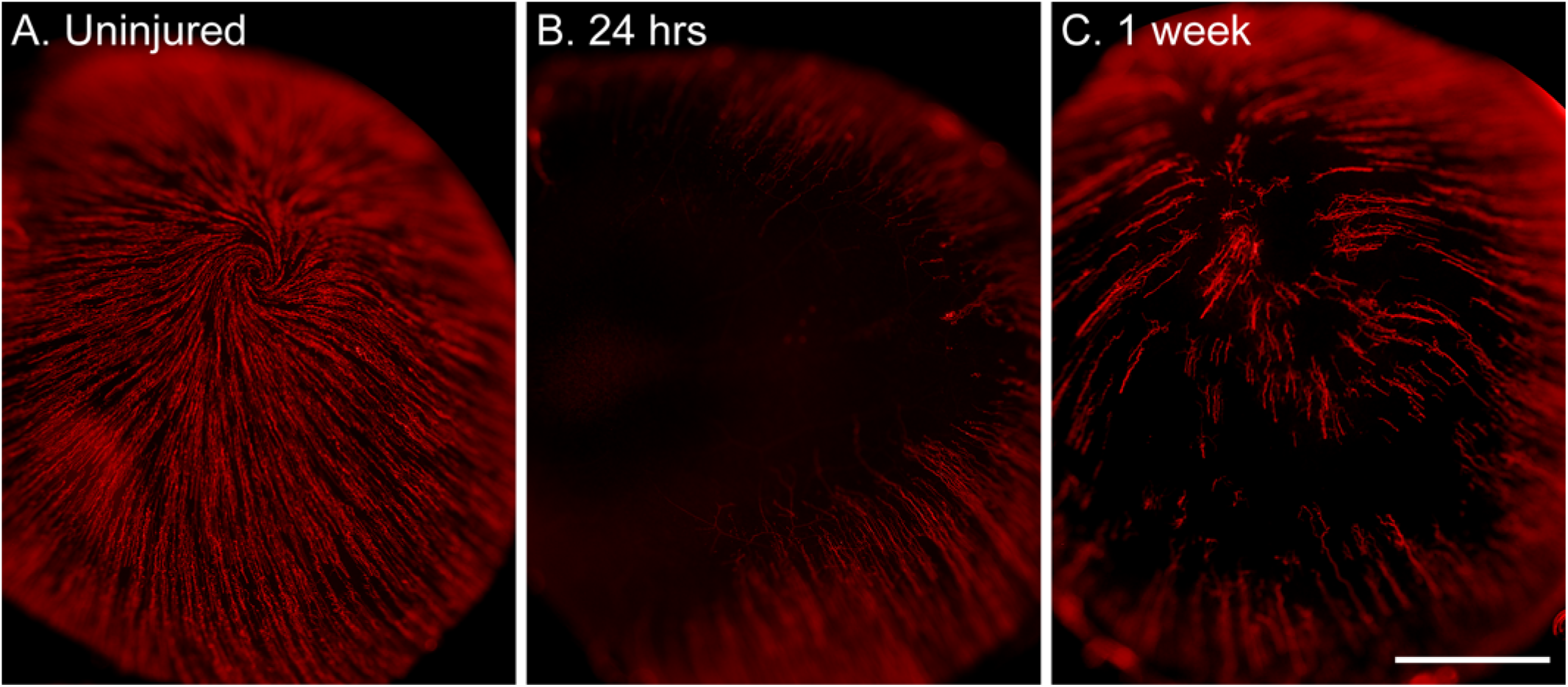
Epifluorescent images of whole mount corneas demonstrate a severe reduction in corneal nerve density in the abraded eye 24 hours after abrasion injury **(B)** compared to an Uninjured cornea **(A)**. There is partial recovery of corneal nerve density one week post-abrasion **(C)**. Scale bar = 1 mm.

Careful examination of nerve endings at the border of the corneal abrasion show enlarged endings resembling neuromas (Fig. 3). Some of these endings contain CGRP immunoreactivity, supporting the idea that these are nociceptors [24]. These endings should be accessible to topical menthol applied to the eye of an awake rat and appear to be sufficient to support hyperalgesia (Fig. 1). Innervation to the central cornea had not returned to control levels one week after injury, and yet, nociceptive responses had returned to baseline levels [50].

**Figure 3.**
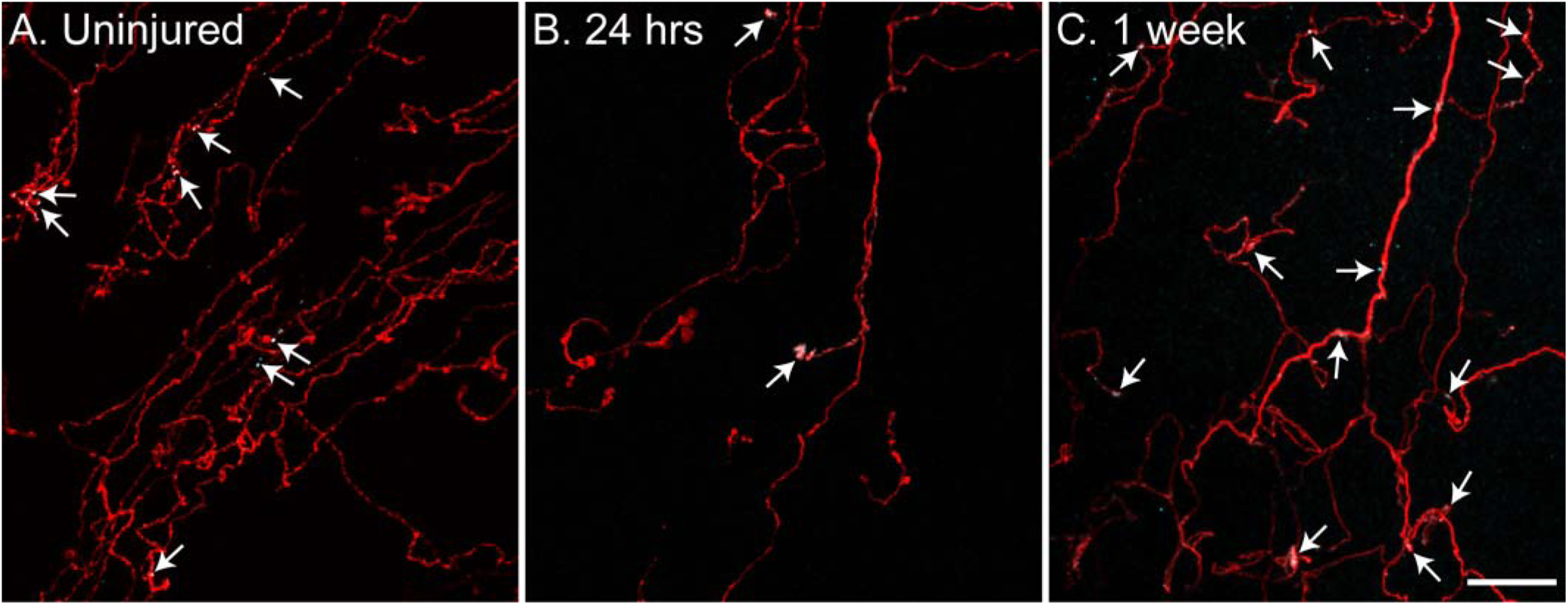
Corneal nerve morphology and CGRP content changes during recovery from abrasion injury. **(A)** A high magnification image of β-tubulin-labeled corneal nerves (red) from an area close to the central whorl of an Uninjured cornea shows thin nerves beaded with varicosities that contain CGRP (blue; arrows). **(B)** At 24 hours post-abrasion, the nerves lining the edge of the abraded area have enlarged nerve endings that contain CGRP (arrows). **(C)** One week after injury, the abraded central cornea is in the process of being repopulated by corneal nerves that appear disorganized and smoother in appearance. CGRP immunoreactivity is found in many of these re-innervating nerves (arrows). Scale bar = 20 μm.

### 3.3. Corneal nerve restoration is produced by original nerve population

Studies of cutaneous nerves suggest that regions of denervated epithelium are re-innervated by new nerve endings taking over the receptive field of the skin that has lost innervation [13]. We tested whether the nerves appearing in the central cornea at 1 week after abrasion injury were from new nerves taking over the corneal receptive field or from the same neurons sending in new branches. This hypothesis was tested by measuring sequential retrograde labeling from the central cornea at the time of abrasion injury (FluoroGold, FG) and again 1 week later (Cholera Toxin B, CTb; Fig. 4A). A smaller application area for the second retrograde dye placement was used to limit labeling to nerve endings within the most central portion of the cornea after injury. Although this approach underestimates the number of cells innervating the central cornea, it ensures that we are only detecting cells with branches in the central cornea. The vast majority (96.2% ± 1.6%, n = 247 neurons from 4 rats) of trigeminal ganglion cells labeled with CTb (Fig. 4B, green) 1 week after abrasion also contained the FG tracer (Fig.4B, magenta) that was applied at the time of injury. These results indicate that the new innervation in the central cornea represents new branches from the nerves that originally occupied this region of the epithelium, not from new nerves taking over that territory.

**Figure 4.**
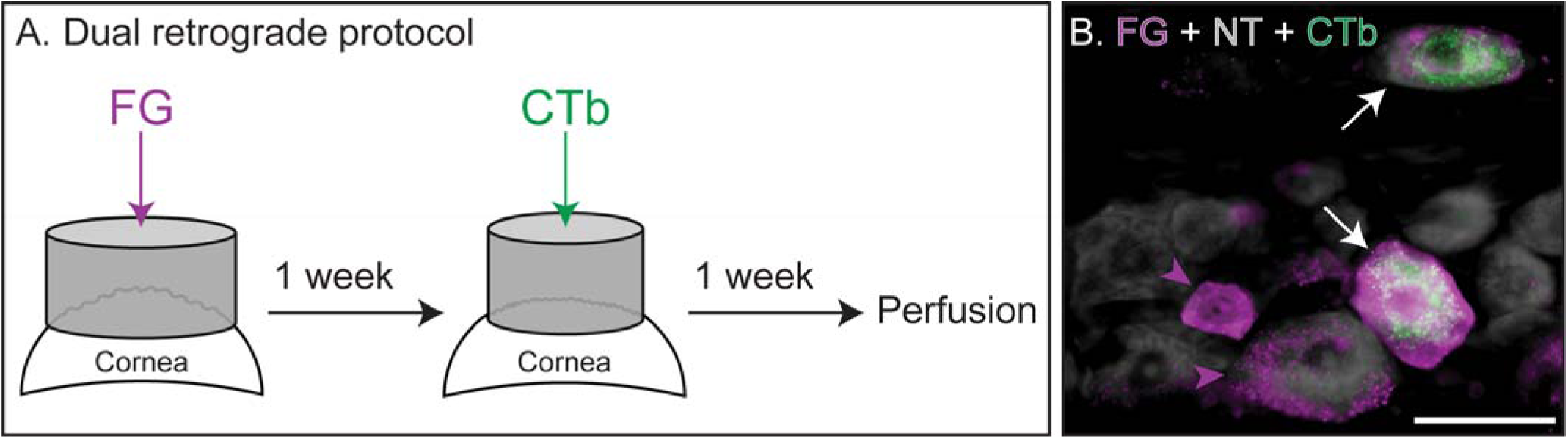
Identification of trigeminal ganglia innervating the central cornea. **(A)** Heptanol was applied within a small 6-gauge metal ring to abrade an area of the central cornea followed by application of the FG tracer. After waiting one week, a shorter heptanol application time within a smaller diameter 7-gauge metal ring was used to lightly abrade the cornea and apply CTb to the central cornea. One week later, trigeminal ganglia were collected for analysis. **(B)** A representative epifluorescent micrograph demonstrates trigeminal ganglia neurons stained with NeuroTrace (NT, gray) that contain only FG (magenta arrowheads), FG and CTb (white arrows) or no tracer. Scale bar = 50 μm.

### 3.4. Unilateral abrasion injury alters corneal nerve density in both corneas

We quantitatively assessed nerve density in the corneal epithelium at different times after injury (Fig. 5). There was a complete absence of corneal nerves in the central portion of the cornea 24 hours after abrasion (Fig. 2B). For the abraded eye, nerve density 1 week after abrasion was reduced by 86% compared to uninjured corneas (Figs. 2, 5; one-way ANOVA (*P* = 0.004), Holm-Sidak post hoc, *P* = 0.006). More surprising was a 56% reduction in corneal nerve volume in the unabraded contralateral eye 24 hours after injury compared to the Uninjured group (Fig. 5; t-test, *P* = 0.0312). The innervation in the contralateral eye returned to baseline levels 1 week after abrasion (Fig. 5; Holm-Sidak post hoc, *P* = 0.956). There were no significant differences in corneal epithelium volume among the treatment groups (Table 2; one-way ANOVA, *P* = 0.358), showing that changes in epithelium volume are not underlying nerve density changes. These findings are noteworthy because there were no detectable changes in nociceptive response in the contralateral eye (Fig. 1) despite significant changes in nerve density.

**Figure 5.**
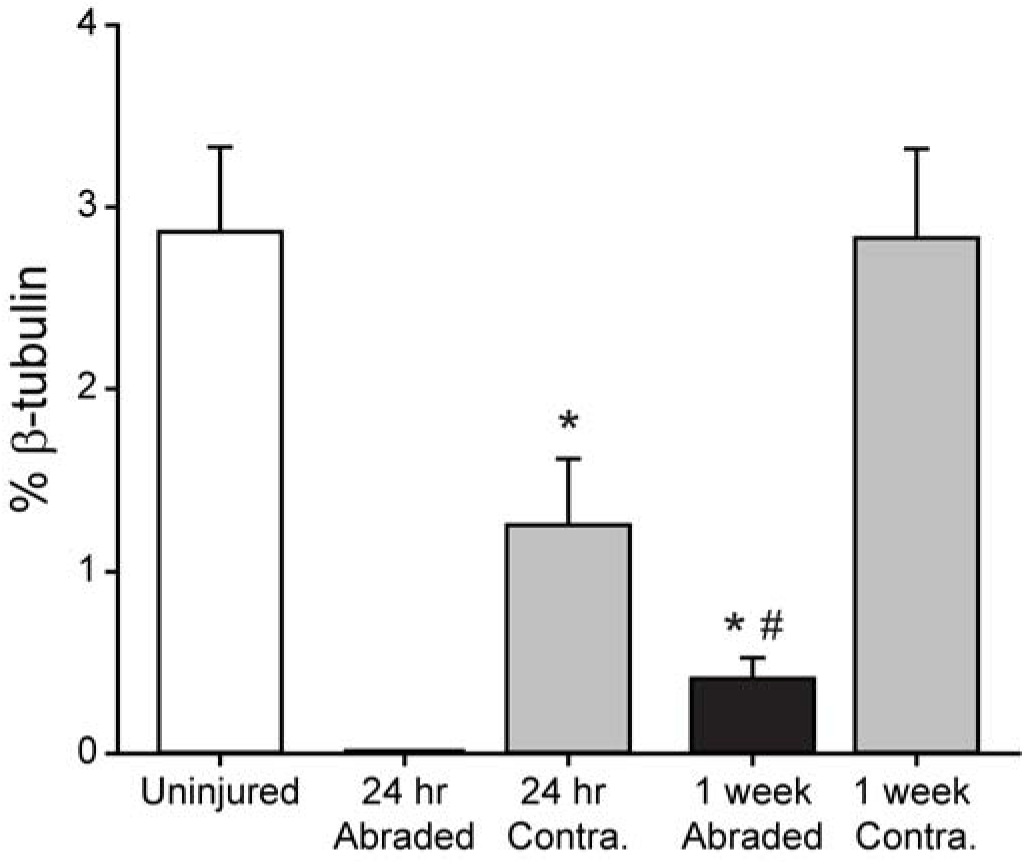
Quantitative changes in corneal epithelial nerve volumes show absence of nerves 24 hours after abrasion in the injured eye, as well as significant nerve loss in the unabraded contralateral eye. Nerve density returned to baseline 1 week later in contralateral, but not abraded corneas. Corneal nerve density was reduced in contralateral corneas 24 hours after abrasion (gray bar, n = 3) compared to uninjured control corneas (white bar, n = 6). Corneal nerve density remains reduced 1 week after abrasion (black bar, n = 4) compared to corneas from uninjured rats and unabraded contralateral corneas 1 week following abrasion (1 week Contra., gray bar, n = 4). * *P* < 0.05 versus Uninjured corneas; # *P* <0.05 versus Contralateral side.

### 3.5. CGRP content is altered after corneal nerve injury

CGRP is the most abundant peptide in corneal central processes terminating in the trigeminal dorsal horn [24], as well as in trigeminal ganglion cell bodies [35; 43]. We examined the epithelial nerves to determine the CGRP content of the most peripheral endings of corneal nerves and how the content of this molecule changes after corneal abrasion injury. CGRP is present in a subset of corneal nerve endings (Fig. 6) and this can be quantified as a percentage of the nerves by examining CGRP as a portion of β-tubulin volume within the epithelium. CGRP could not be calculated in the abraded corneas 24 hours after injury because intraepithelial nerves were absent, but 1 week after abrasion the CGRP content of the remaining nerves was increased approximately four-fold compared to the Uninjured control group (Fig. 7; Kruskal-Wallis one-way ANOVA on Ranks, *P* = 0.003; Dunn’s Method post hoc test, *P* < 0.05). Interestingly, CGRP content of nerves in the contralateral eye was stable at both 24 hour and 1 week time points, despite significant fluctuations in nerve volume in the uninjured eye (Fig. 5). This suggests that the neuropeptide content is well regulated during changes in innervation density in the absence of injury to the cornea. This contrasts with the injured cornea where CGRP content was elevated even when nerve density was quite low (Fig. 6, 7). It is also noteworthy that CGRP immunoreactivity was present in abraded corneas 1 week after injury in a diffuse pattern (Fig. 6E) that was not associated with nerve endings, and therefore not included in our analysis, but could be interpreted as being present in epithelial tissue. We speculate that this could be due to increased peptide release from the corneal nerves. Our findings support the hypothesis that there are phenotypic changes in corneal nerves that occur specifically after peripheral nerve axotomy.

**Figure 6.**
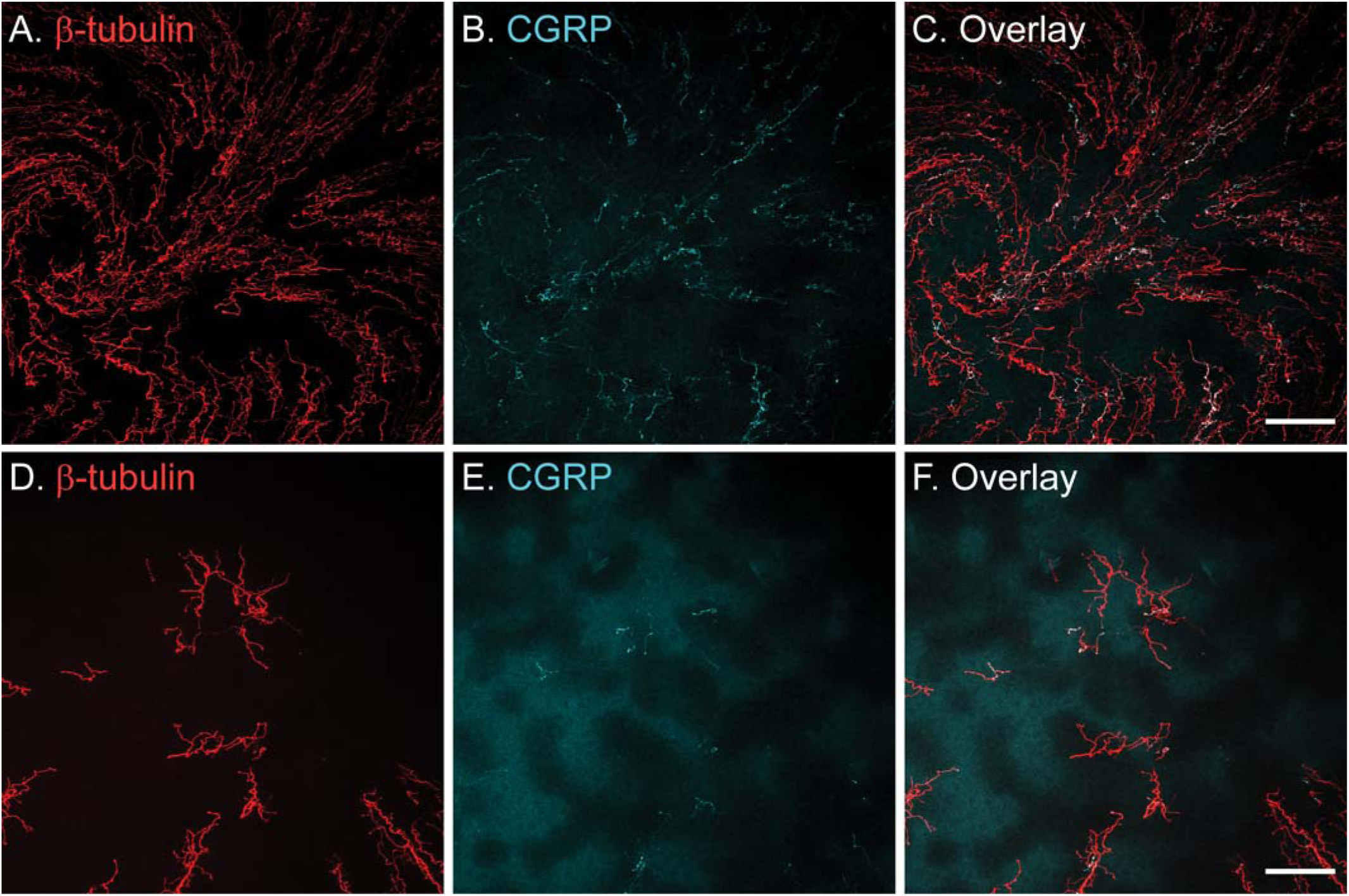
Confocal images of central corneas demonstrate partial recovery of β-tubulin-ir nerves at 1 week post-abrasion (**D – F**) as compared to an uninjured control cornea (**A – C**). However, β-tubulin **(D)** and, as a result, CGRP **(E)** expression remains low at 1 week post-abrasion **(E)**. Scale bar = 100 μm.

**Figure 7.**
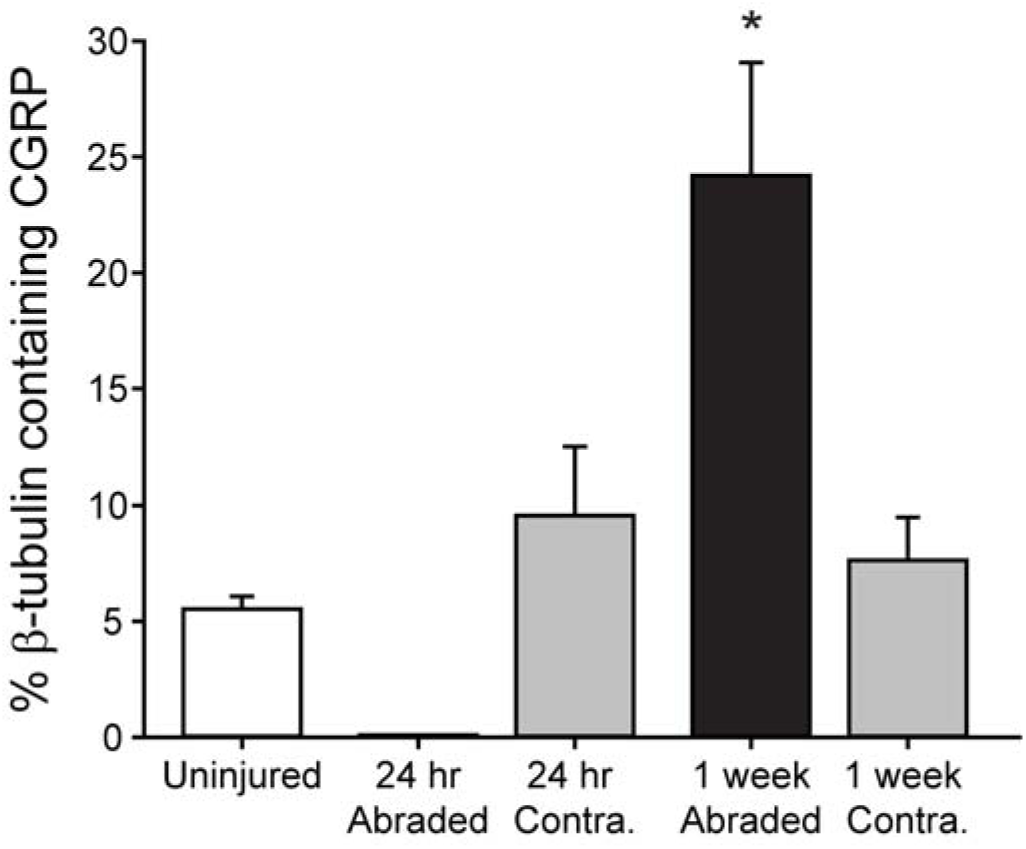
CGRP content of β-tubulin nerves was increased 1 week after abrasion in injured corneas, but not in the nerves of the contralateral eye despite reductions in nerve density (compare with Fig. 5). The volume of CGRP immunoreactivity within the volume of β-tubulin-ir corneal nerves was calculated for each cornea and expressed as the percent of β-tubulin containing CGRP. CGRP content was not measurable at 24 hours in the abraded corneas because intraepithelial nerves were absent in these animals. The percent of β-tubulin volume containing CGRP was increased 1 week after abrasion injury (black bar, n = 4) as compared to Uninjured corneas (white bar, n = 6). CGRP levels remained stable within contralateral corneas at 24 hours (gray bar, n = 3) and 1 week (gray bar, n = 4) post-abrasion. * *P* < 0.05 compared to the Uninjured group.

### 3.6. Corneal abrasion causes delayed reduction in tear production

Corneal abrasions can evoke symptoms of dry eye, including changes in tear production [2]. Tear production in the abraded eye was significantly reduced one week after the abrasion (Baseline: 13.5 ± 1.5 mm; 1 week Abraded: 9.0 ± 1.3 mm; Paired t-test, *P* = 0.0489), but not 24 hours after abrasion (Baseline: 12.8 ± 1.4; 24 hrs Abraded: 13.9 ± 1.5; Paired t-test, *P* = 0.182). The tear measurements in the contralateral eye were not different at 24 hours (Baseline: 9.5 ± 1.6; 24 hrs Contralateral: 9.5 ± 1.4; Paired t-test, P = 0.5) or at 1 week (Baseline: 8.5 ± 1.0; 1 week Contralateral: 7.0 ± 1.1; Paired t-test, P = 0.148), suggesting some independence of the reflex circuits for the two eyes. The percent of baseline tear production was calculated for each animal, averaged for each treatment group and expressed as % Tears (Fig. 8). The temporal expression of the changes in tear production are apparently unrelated to either nociceptive responses (Fig. 1) or corneal nerve density (Fig. 5). Tear production was normal 24 hours after abrasion in both eyes when epithelial nerve density was absent or significantly reduced, but was reduced 1 week after abrasion when innervation has begun to be restored (Fig. 8). Interestingly, this is a time when expression of CGRP is elevated in the nerves (Fig. 7), which may impact epithelial health.

**Figure 8.**
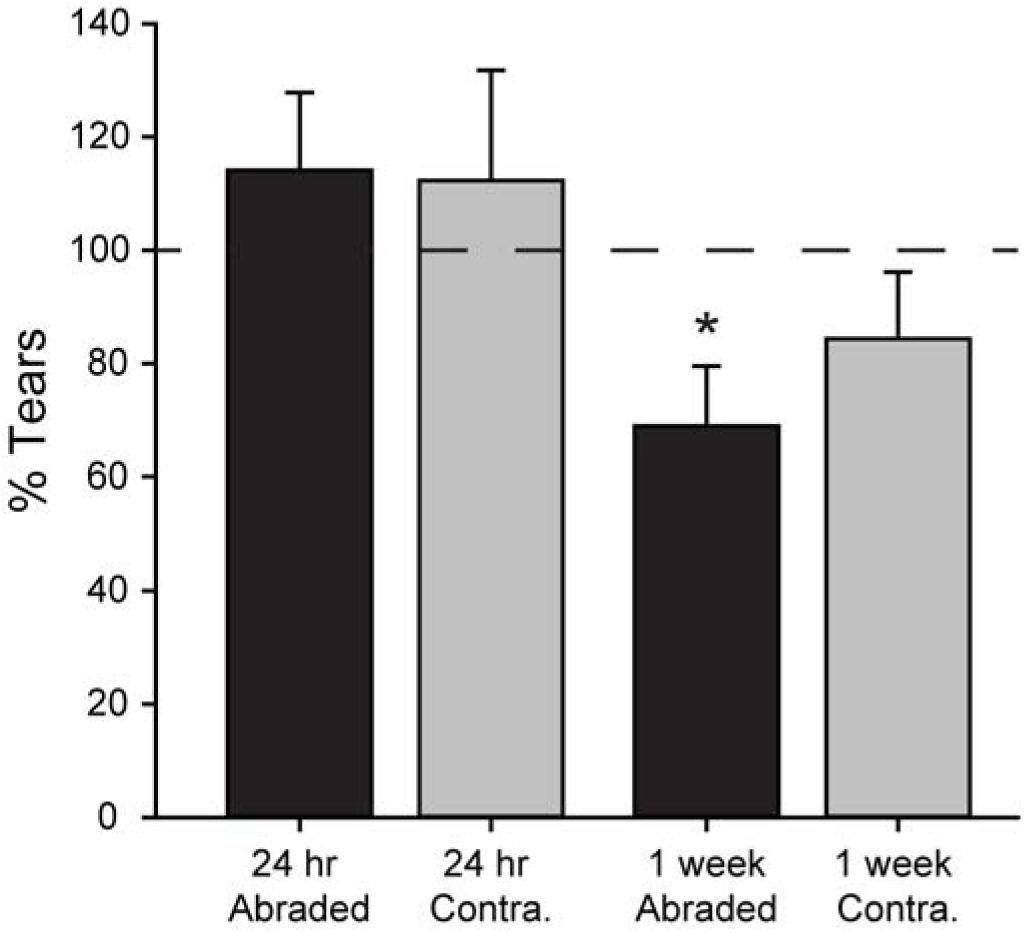
Abrasion injury has a delayed effect on tear production. Phenol thread measurements taken at 24 hours or 1 week after injury are expressed as a percent of pre-injury baseline measurements (% Tears; Baseline: dotted line at 100%). Tear production was not affected 24 hours after abrasion injury on either the injured (24 hr Abraded, black bar, n = 8) or unabraded (24 hr Contralateral, gray bar, n = 8) side. Tear production was significantly reduced 1 week after abrasion injury on the injured side (1 week Abraded, black bar, n = 4) as compared to pre-injury, but not the unabraded (1 week Contralateral, gray bar, n = 4) side. * *P* < 0.05 as compared to raw pre-injury phenol thread measurements.

### 3.7. Corneal abrasion increases unilateral eye closures (winks) but not blinks

We assessed changes in behavioral responses related to ocular surface homeostasis since molecular mechanisms involved in cold nociception [46] have also been implicated in both spontaneous blinking and tear production [45; 48]. We assessed bilateral (blinks; Fig. 9A) and unilateral (winks; Fig. 9B) eye closures in subsets of our corneal injury rats and uninjured control rats. We were not able to accurately quantify blinking and winking behavior at 24 hours after injury because these rats exhibit profound photophobia including extended eye closures [19]. However, there were increased ipsilateral winks one week after abrasion compared to Baseline (Fig. 9B; Paired t-test, *P* = 0.0009). Abraded animals also demonstrated increased winks in the contralateral unabraded eye (Fig. 9B; Paired t-test, *P* = 0.01), although bilateral blink frequency did not change significantly (Fig. 9A; Paired t-test, *P* = 0.324). These results show that the overall rate of eye closure is increased, but the responses are not bilaterally coordinated. This change in eye closure behavior occurs at a time when nocifensive responses have returned to normal, but innervation is still significantly reduced (Fig. 5) and tear production is also reduced (Fig. 8). Interestingly, the increased wink behavior was detected in both eyes, while tear production was only significantly reduced in the abraded eye.

**Figure 9.**
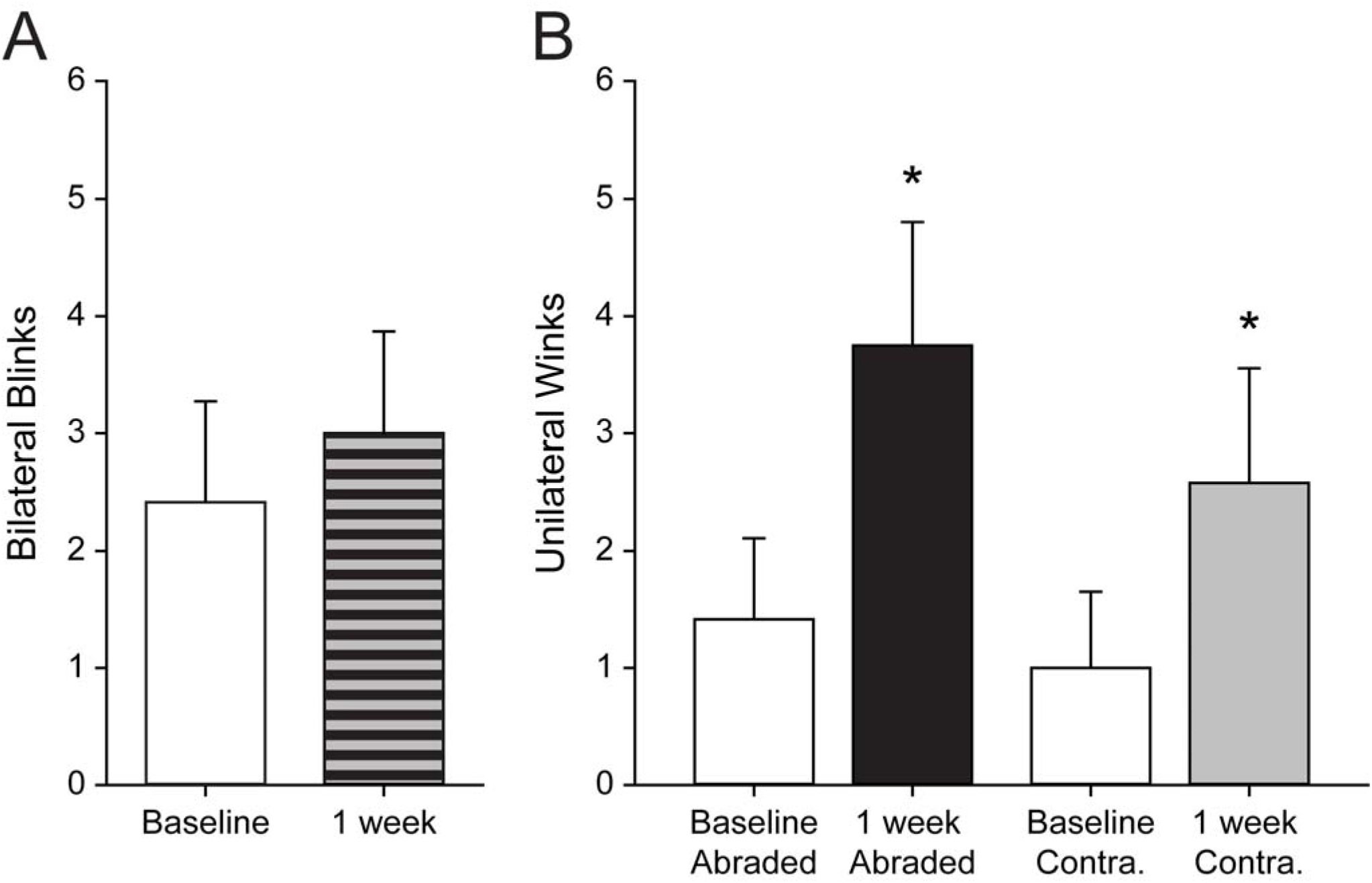
Eye closures were differentially affected by abrasion injury. Paired t-tests were run between the number of bilateral blinks or unilateral winks assessed before injury (Baseline) and 1 week after injury. It was not possible to assess blinks and winks at 24 hours post-abrasion due to photophobia which produced prolonged eye closures. **(A)** Spontaneous bilateral blinks were not different 1 week after abrasion injury (black/gray patterned bar, n = 12) as compared to pre-injury Baseline assessments (white bar, n = 12). **(B)** Unilateral winks increased 1 week after abrasion in both the abraded (black bar, n = 12) and contralateral eyes (gray bar, n = 12) as compared to pre-injury Baseline (white bars). * *P* < 0.05 as compared to Baseline.

### 3.8. Increased expression of ATF3 and CGRP in trigeminal ganglia after injury

Our findings show a large disconnect between corneal epithelial nerve density, pain responses, and homeostatic responses such as tear production and eye closure. We hypothesize that phenotypic changes in corneal nerve cell bodies are likely induced after frank injury of the peripheral endings in the epithelium. If this is the case, we would expect to see changes in the expression of molecules in the trigeminal ganglion that happen acutely after injury. We examined the expression of a neuronal injury marker, ATF3 [52], as well as CGRP, the primary neuropeptide in corneal nociceptors [24]. Both ATF3 and CGRP are present in subsets of trigeminal ganglion cells after injury (Fig. 10). The proportion of trigeminal ganglion cells containing the nerve injury marker ATF3 was increased 24 hours after injury in Abraded rats, but only on the injured side (Fig. 11A) compared to Uninjured control animals (one-way ANOVA (P = 0.007), Holm-Sidak post hoc, *P* = 0.008) or the contralateral unabraded side (Holm-Sidak post hoc, *P* = 0.027). ATF3 levels returned to control levels 1 week post-abrasion (one-way ANOVA, *P* = 0.343). The nociceptive marker CGRP showed the same pattern of change in the trigeminal ganglion, with increased levels 24 hours after injury on the side ipsilateral to the abrasion (Fig. 11B; one-way ANOVA, *P* = 0.046). Interestingly, the CGRP content of trigeminal ganglion cells returned to baseline levels 1 week after injury (one-way ANOVA, *P* = 0.322) when CGRP content in the peripheral endings was increased (Fig. 7). Thus, the elevated CGRP content in the trigeminal ganglion is better correlated with increased pain responses than the CGRP content in the peripheral endings within the cornea, at the 1 week time point when both portions of the trigeminal nerve were assessed. Despite changes in peripheral nerve density in the contralateral eye 24 hours after injury (Fig. 5), there was no change in expression of either ATF3 or CGRP on the contralateral side at either time point. This difference suggests that the changes in peripheral nerve density seen in the contralateral eye is not of sufficient degree or nature to evoke molecular changes in the trigeminal ganglion, the cell bodies of the peripheral endings in the cornea.

**Figure 10.**
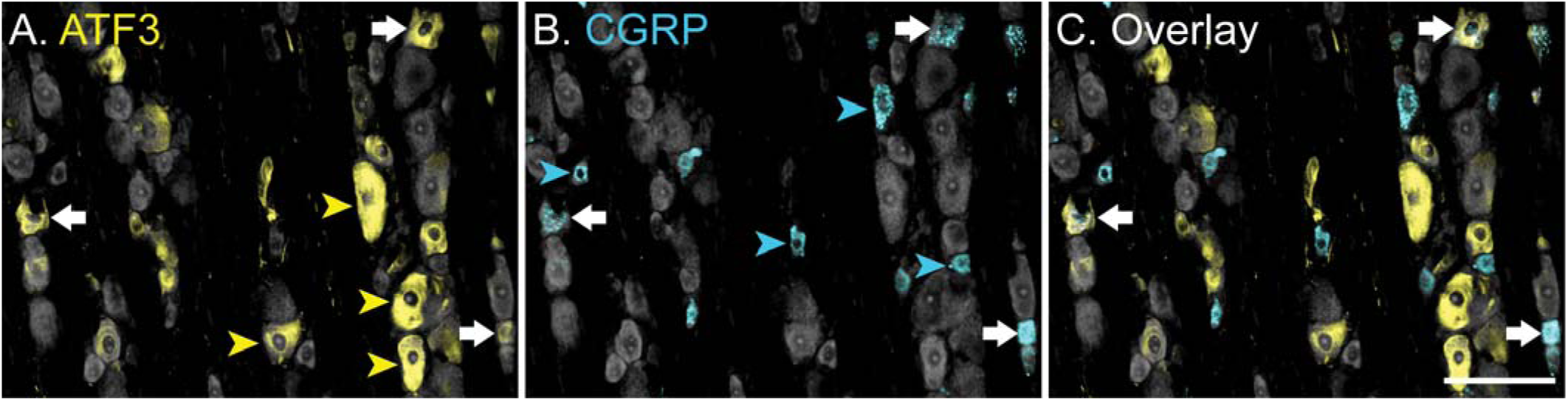
ATF3, a marker of neuronal injury, and the nociceptive marker CGRP are both present in the rat trigeminal ganglia. Subsets of trigeminal ganglion neurons (gray) contain ATF3 (**A**, yellow arrowheads) or CGRP (**B**, blue arrowheads), or in some cases, both markers (**C**, white arrows). Scale bar = 50 μm.

**Figure 11.**
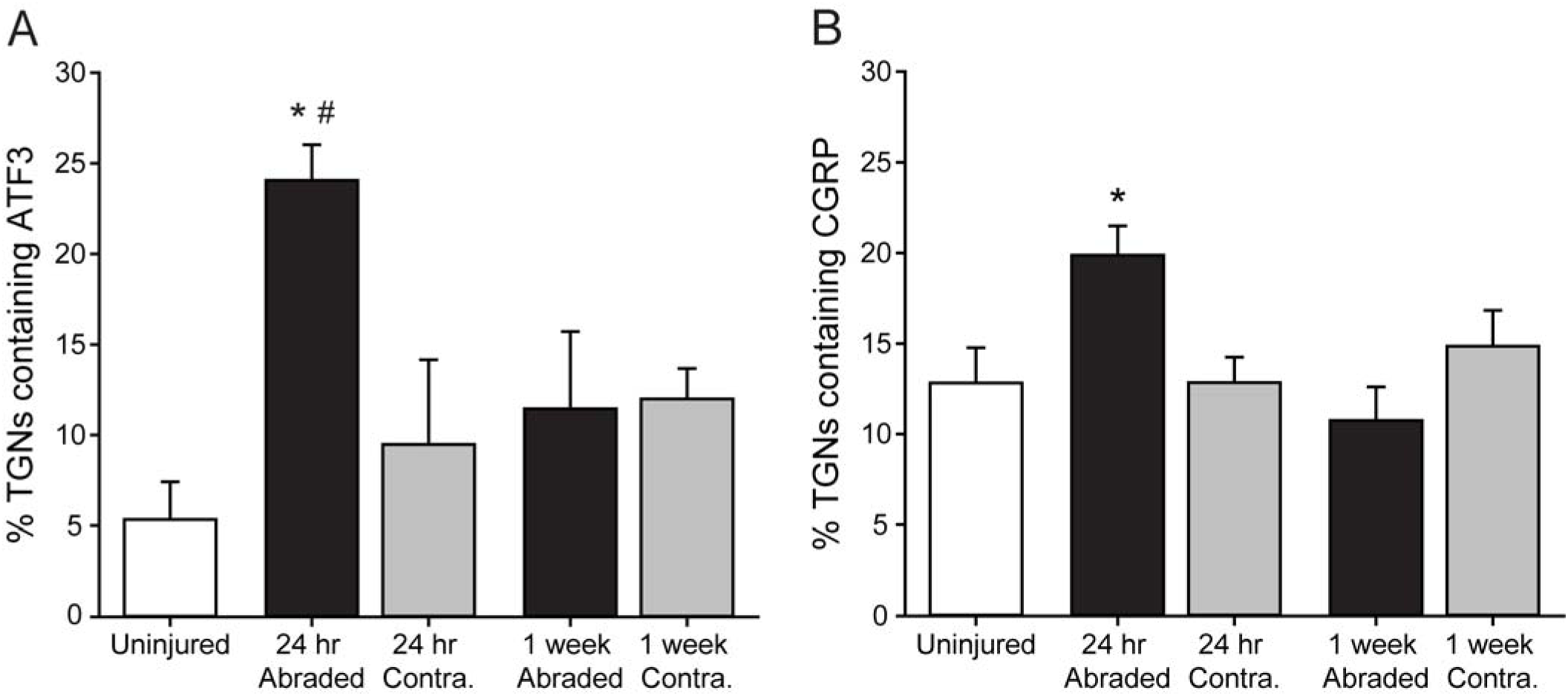
ATF3 and CGRP are increased in trigeminal ganglion neurons (TGNs) after abrasion injury. The percent of TGNs that contain (**A**) ATF3 or (**B**) CGRP is increased in ipsilateral TGNs 24 hours after corneal abrasion (black bars, n = 3) compared to Uninjured rats (white bars, n = 4). One week post-abrasion the percent of TGNs on the abraded side containing ATF3 (**A**) or CGRP (**B**) is back to control levels (black bars, n = 6). There were no significant increases in ATF3 or CGRP in contralateral TGNs at either time point (24 hr, gray bars, n = 3; 1 week, gray bars, n = 6). * *P* < 0.05 versus Uninjured; # *P* <0.05 versus Contralateral side.

## 4. Discussion

Our findings show complex changes in pain, corneal nerve density, molecular expression, and ocular homeostasis after a superficial abrasion of the cornea. Our findings support the idea that corneal nerve density per se is not a good predictor of pain state or ocular homeostasis. Pain responses occur acutely and are best correlated with increased expression of ATF3 and CGRP in the trigeminal ganglion; while dry eye symptoms, including increased eye closures and reduced tear production, occur at later time points when acute pain has resolved.

A chemical abrasion of the corneal surface with heptanol virtually abolished the epithelial innervation, but resulted in increased pain responses. This paradoxical finding [50] suggests that abrasion injury causes alterations in either nerve function or the central pain pathways mediating corneal sensation. The presence of neuromas at the edge of the abrasion injury may be a local mechanism for increased sensory nerve activation. We found expression of CGRP in these neuromas and others have found accumulation of sodium channels at these sites, coupled with increase excitability of injured nerves [3; 11; 12; 40]. Blockade of sodium channels with TTX prevents the photophobia seen with corneal heptanol injury [19]. Since both the light used to evoke photophobia and our topical menthol application are able to reach nerve endings at the edge of the corneal injury, our findings are consistent with an acute upregulation of sodium channels in these endings that may mediate hyperalgesia. Changes in corneal nerve morphology, including swellings, have been reported after both nerve injury and application of intense stimuli such as high osmotic concentrations of hypertonic saline [23; 28] suggesting that these enlargements may be a site for aberrant nerve activity. In contrast, responses to stimuli confined to the denervated central cornea are impaired in the hours immediately after heptanol abrasion [27].

Both CGRP and ATF3 expression in trigeminal ganglia cell bodies increased 24 hours after injury and returned to normal levels by 1 week, paralleling the time course of changes in nociceptive responses. The increased expression of CGRP and ATF3 in the trigeminal ganglion may enhance central activation of pain pathways in response to ocular stimulation, leading to hyperalgesia [36]. Surprisingly, the CGRP content of the trigeminal ganglion was normal one week after injury, while CGRP expression was elevated within the peripheral corneal nerve population that is re-innervating the central cornea. In a prior study, no changes in CGRP or Substance P in trigeminal ganglion cells were seen after corneal injury with sodium hydroxide (NaOH) when assessed three days after injury [16]. Our findings suggest that changes in molecular expression within the trigeminal ganglia are rapid and dynamic after injury. A previous study from our laboratory found expression of the Transient Receptor Potential Vanilloid 1 (TRPV1) channel in 30% of corneal-projecting cells in the trigeminal ganglia, but only 2% of their central terminals [22], consistent with the idea of differential trafficking of molecules within corneal sensory neurons.

Increased CGRP expression in corneal epithelial nerves was seen in the context of severely reduced corneal nerve density, suggesting that there is more CGRP per nerve ending in the ocular surface. CGRP is known to be released upon noxious corneal stimulation [3] and we can speculate that there may be increased CGRP release in the epithelium despite reduced nerve density. Reduced number of nerves and increased release of CGRP may impair corneal epithelial function and eventually destabilize the corneal surface [33]. One long-term consequence of this impaired corneal epithelial function may be reduced tear production [41], and the reduced tear production may stimulate reflex blinking [25], and thus cause the increase in eye closure responses at 1 week after injury.

We saw no evidence of altered nociceptive responses in the contralateral eye, despite a 56% reduction in corneal nerve density [34]. Prior studies have reported similar findings in which a unilateral disease or injury in the cornea will produce bilateral effects on corneal nerve density, with no changes in corneal sensation. Laser *in vivo* confocal microscopy is used to assess corneal sub-basal nerve density in humans [54]. Patients with unilateral microbial keratitis had significant decreases in total number of corneal nerves, total nerve length, number of branches and branch nerve length in the uninfected, contralateral cornea [10]. Despite this decrease, there was no significant change in corneal sensation. In a mouse model, after unilateral trigeminal axotomy sub-basal nerves from the center of the contralateral cornea were reduced one day after axotomy, which caused a loss of sensation on the injured side; on the contralateral side, there was a loss of nerves, but no significant decrease in corneal sensation [57]. The mechanisms underlying these contralateral changes are unclear, especially given that there are other examples of unilateral corneal injury and disease states that affect both contralateral nerve density and sensation [21]. Possible mechanisms of bilateral changes induced by unilateral corneal injury include bilateral increases in dendritic cells which mediate immune responses and alterations in tear cytokines in the context of unilateral bacterial keratitis [56]. Others have suggested that contralateral hyperalgesia, or mirror pain, after nerve injury may be related to changes in the peripheral nervous system, central pathways and/or the activation of immunocompetent cells such as cytokines, macrophages, microglia and astrocytes [15; 49]. The presence of contralateral sensory deficits have also been reported in various genetic knockout animals, included VIP and TNF alpha receptor knockouts [17; 38]. Together, these studies suggest that the causes of contralateral effects of nerve injury may stem from multiple mechanisms, and may depend on the type and severity of the injury or disease state.

Spontaneous pain was evident as a depression in home cage wheel running 24 hours after corneal injury. Previous studies have shown that home cage wheel running is a sensitive method to objectively assess the duration and magnitude of inflammatory pain and migraine-like pain [31; 32]. Despite the extent of the nerve injury, the pain resolved quite rapidly with wheel running returning to baseline levels quickly after injury; supporting the argument that pain is largely resolved within a few days despite the lingering loss of corneal epithelial innervation.

Our animals also exhibited extensive blinking and photophobia 24 hours after abrasion. At one week post-abrasion, bilateral blinks were at baseline levels while unilateral eye closures were increased in both the injured and contralateral eye. We speculate that the bilateral blinks may represent the spontaneous blink pattern [30] in these animals while the unilateral eye closures, or winks, may represent corneal reflexes from ongoing spontaneous pain after abrasion injury. Previous studies in humans looking at normal blink rates and blink patterns found that even after topical application of an anesthetic to the cornea, spontaneous blinking was reduced but still present [44]. These and other studies suggest that spontaneous blinks are centrally-mediated and modulated by external factors [5; 30], rather than being solely dependent on the state of the peripheral corneal environment. A recent human study found a correlation between higher blink rates and lower thresholds to corneal stimulation, meaning that more corneal pain is correlated with higher rates of blinking [44]. Therefore, if the rat is experiencing bouts of spontaneous pain, this may increase unilateral corneal reflexive eye closures on the injured side. It is unclear why the unabraded eye experienced increased unilateral eye closures as well, but causes of contralateral effects of injury or disease are complicated and require further study.

Corneal nerves are critically important in maintaining the health and integrity of the corneal epithelial environment [6; 42]. In light of the reduction in corneal nerve density, we would have predicted that abrasion injury would affect tear production at both time points post-injury. However, we only found a reduction in tears at the one week time point. It is possible that the extensive blinking observed at 24 hours may have produced reflex tears that would have masked a decrease in normal tear production caused by corneal nerve damage. At one week, the decrease in tears may indicate subtle, persistent effects of corneal nerve injury. Our study has demonstrated that the cornea is a complex system in which nociceptive and homeostatic mechanisms are integrated in order to maintain the important protective barrier. A recent study in humans demonstrated that corneal sensation, tear film stability, tear break-up time, and blink rate are significantly correlated [44]. This integration may underlie the temporal dynamics of corneal injury (Fig. 12) with immediate paradoxical changes in nerve density and evoked corneal sensation giving way to a dry eye disease phenotype with changes in spontaneous pain, blink rate, and tear production. This study also suggests that corneal nerve density alone is not a good predictor of pain state [18].

**Figure 12.**
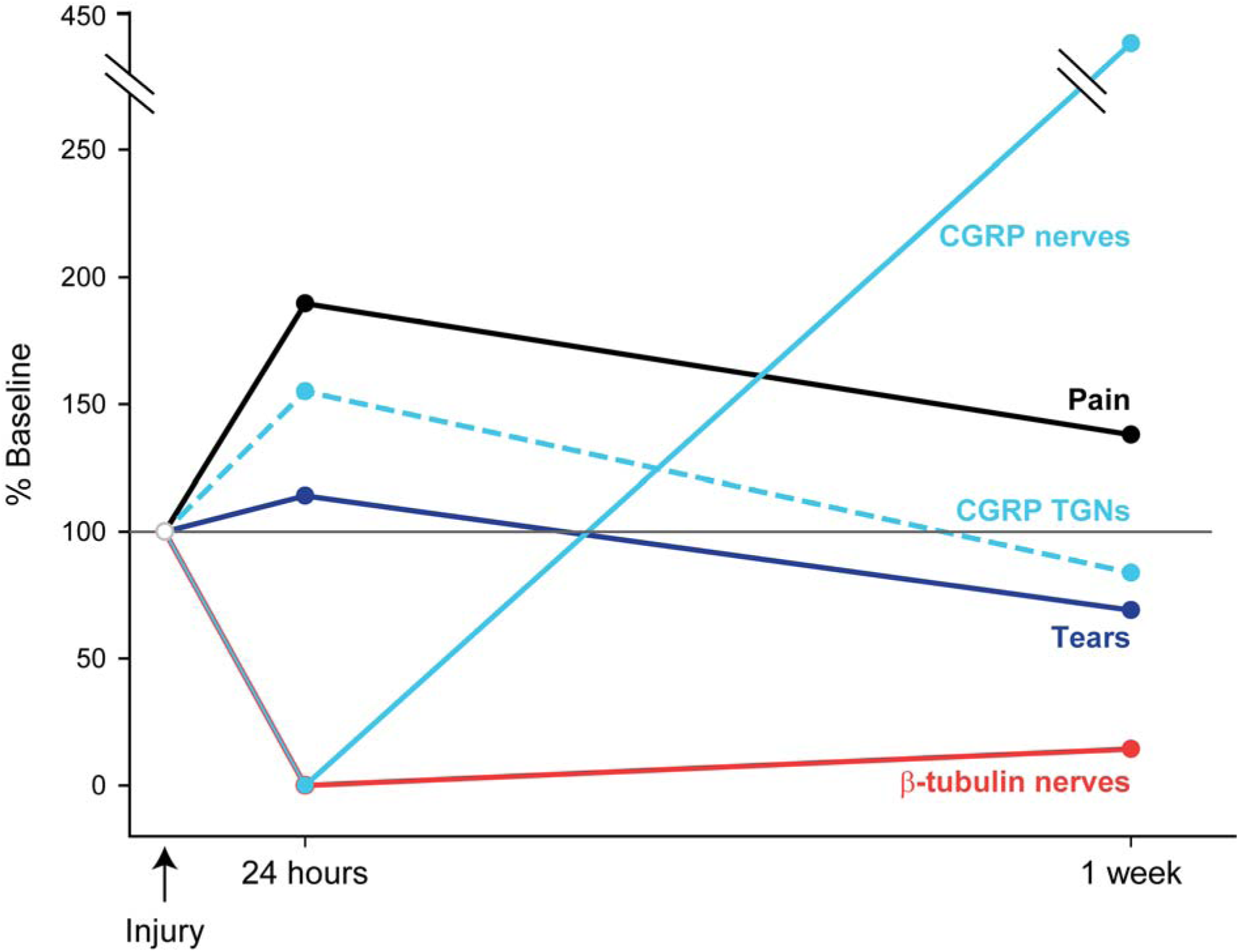
Corneal injury alters nociceptive and homeostatic responses with distinct temporal dynamics. All data points represent the percent of baseline values of the abraded side of injured animals; a simple straight line graph was generated using X (time) and Y (% Baseline) pairs of data points within each metric. The graph is not intended to assume a linear trend between 24 hours and 1 week, but illustrates the direction of post-abrasion changes. Paradoxically, as corneal nerve density in the central cornea was eliminated (red line, 24 hours), injured animals demonstrated hyperalgesia to noxious menthol stimulation (black line, 24 hours). Tear production was decreased one week after abrasion injury (blue line, 1 week), but not acutely (blue line, 24 hours) on the side ipsilateral to injury. There is an increase in the percent of β-tubulin-ir corneal nerves that contain CGRP at 1 week post-injury (CGRP nerves, solid cyan line, 1 week). The number of trigeminal ganglion neurons containing CGRP (CGRP TGNs, dashed cyan line) increases at 24 hours, mirroring the hyperalgesia measured in abraded animals, and then decreases by 1 week post-injury.

## Acknowledgments

The authors wish to thank Emma Eikerman, Aveek Ganguly, Clayton Hudson, and Ram Kandasamy for their technical assistance. The work was supported by grants from NIH: P30 NS061800 and NS095057; the Medical Research Foundation of Oregon; and Oregon Health & Science University.

## References

[1] Aicher SA, Hermes SM, Hegarty DM. Denervation of the Lacrimal Gland Leads to Corneal Hypoalgesia in a Novel Rat Model of Aqueous Dry Eye Disease. Investigative ophthalmology & visual science 2015;56(11):6981–6989.

[2] Ang RT, Dartt DA, Tsubota K. Dry eye after refractive surgery. CurrOpinOphthalmol 2001;12(4):318–322.

[3] Belmonte C, Acosta MC, Gallar J. Neural basis of sensation in intact and injured corneas. ExpEye Res 2004;78(3):513–525.

[4] Belmonte C, Brock JA, Viana F. Converting cold into pain. Exp Brain Res 2009;196(1):13–30.

[5] Belmonte C, Nichols JJ, Cox SM, Brock JA, Begley CG, Bereiter DA, Dartt DA, Galor A, Hamrah P, Ivanusic JJ, Jacobs DS, McNamara NA, Rosenblatt MI, Stapleton F, Wolffsohn JS. TFOS DEWS II pain and sensation report. The ocular surface 2017;15(3):404–437.

[6] Beuerman RW, Schimmelpfennig B. Sensory denervation of the rabbit cornea affects epithelial properties. Exp Neurol 1980;69(1):196–201.

[7] Braz JM, Basbaum AI. Differential ATF3 expression in dorsal root ganglion neurons reveals the profile of primary afferents engaged by diverse noxious chemical stimuli. Pain 2010;150(2):290–301.

[8] Carlton SM, Westlund KN, Zhang D, Sorkin LS, Willis WD. Calcitonin gene-related peptide containing primary afferent fibers synapse on primate spinothalamic tract cells. Neuroscience Letters 1990;109:76–81.

[9] Chao C, Golebiowski B, Stapleton F. The role of corneal innervation in LASIK-induced neuropathic dry eye. The ocular surface 2014;12(1):32–45.

[10] Cruzat A, Schrems WA, Schrems-Hoesl LM, Cavalcanti BM, Baniasadi N, Witkin D, Pavan-Langston D, Dana R, Hamrah P. Contralateral Clinically Unaffected Eyes of Patients With Unilateral Infectious Keratitis Demonstrate a Sympathetic Immune Response. Investigative ophthalmology & visual science 2015;56(11):6612–6620.

[11] Davies SL, Loescher AR, Clayton NM, Bountra C, Robinson PP, Boissonade FM. Changes in sodium channel expression following trigeminal nerve injury. Exp Neurol 2006;202(1):207–216.

[12] Devor M, Keller CH, Deerinck TJ, Levinson SR, Ellisman MH. Na+ channel accumulation on axolemma of afferent endings in nerve end neuromas in Apteronotus. Neurosci Lett 1989;102(2–3):149–154.

[13] Devor M, Schonfeld D, Seltzer Z, Wall PD. Two modes of cutaneous reinnervation following peripheral nerve injury. J Comp Neurol 1979;185(1):211–220.

[14] Dougherty PM, Willis WD. Modification of the responses of primate spinothalamic neurons to mechanical stimulation by excitatory amino acids and an N-methyl-D-aspartate antagonist. Brain Res 1991;542:15–22.

[15] Dubovy P, Tuckova L, Jancalek R, Svizenska I, Klusakova I. Increased invasion of ED-1 positive macrophages in both ipsi- and contralateral dorsal root ganglia following unilateral nerve injuries. Neurosci Lett 2007;427(2):88–93.

[16] Felipe CD, Gonzalez GG, Gallar J, Belmonte C. Quantification and immunocytochemical characteristics of trigeminal ganglion neurons projecting to the cornea: effect of corneal wounding. EurJ Pain 1999;3(1):31–39.

[17] Gallo A, Leerink M, Michot B, Ahmed E, Forget P, Mouraux A, Hermans E, Deumens R. Bilateral tactile hypersensitivity and neuroimmune responses after spared nerve injury in mice lacking vasoactive intestinal peptide. Exp Neurol 2017;293:62–73.

[18] Galor A, Levitt RC, Felix ER, Martin ER, Sarantopoulos CD. Neuropathic ocular pain: an important yet underevaluated feature of dry eye. Eye (Lond) 2015;29(3):301–312.

[19] Green PG, Alvarez P, Levine JD. Topical Tetrodotoxin Attenuates Photophobia Induced by Corneal Injury in the Rat. The journal of pain: official journal of the American Pain Society 2015;16(9):881–886.

[20] Grixti A, Sadri M, Watts MT. Corneal protection during general anesthesia for nonocular surgery. The ocular surface 2013;11(2):109–118.

[21] Hamrah P, Cruzat A, Dastjerdi MH, Pruss H, Zheng L, Shahatit BM, Bayhan HA, Dana R, Pavan-Langston D. Unilateral herpes zoster ophthalmicus results in bilateral corneal nerve alteration: an in vivo confocal microscopy study. Ophthalmology 2013;120(1):40–47.

[22] Hegarty DM, Hermes SM, Largent-Milnes TM, Aicher SA. Capsaicin-responsive corneal afferents do not contain TRPV1 at their central terminals in trigeminal nucleus caudalis in rats. J Chem Neuroanat 2014;61–62C:1–12.

[23] Hegarty DM, Hermes SM, Yang K, Aicher SA. Select noxious stimuli induce changes on corneal nerve morphology. J Comp Neurol 2017;525(8):2019–2031.

[24] Hegarty DM, Tonsfeldt K, Hermes SM, Helfand H, Aicher SA. Differential localization of vesicular glutamate transporters and peptides in corneal afferents to trigeminal nucleus caudalis. J Comp Neurol 2010;518(17):3557–3569.

[25] Henriquez VM, Evinger C. The three-neuron corneal reflex circuit and modulation of second-order corneal responsive neurons. Experimental Brain Research 2007;179(4):691–702.

[26] Hermes SM, Andresen MC, Aicher SA. Localization of TRPV1 and P2X3 in unmyelinated and myelinated vagal afferents in the rat. J Chem Neuroanat 2016;72:1–7.

[27] Hirata H, Mizerska K, Dallacasagrande V, Guaiquil VH, Rosenblatt MI. Acute corneal epithelial debridement unmasks the corneal stromal nerve responses to ocular stimulation in rats: implications for abnormal sensations of the eye. J Neurophysiol 2017;117(5):1935–1947.

[28] Hirata H, Mizerska K, Marfurt CF, Rosenblatt MI. Hyperosmolar Tears Induce Functional and Structural Alterations of Corneal Nerves: Electrophysiological and Anatomical Evidence Toward Neurotoxicity. Investigative ophthalmology & visual science 2015;56(13):8125–8140.

[29] Kallinikos P, Berhanu M, O’Donnell C, Boulton AJ, Efron N, Malik RA. Corneal nerve tortuosity in diabetic patients with neuropathy. Invest OphthalmolVisSci 2004;45(2):418–422.

[30] Kaminer J, Powers AS, Horn KG, Hui C, Evinger C. Characterizing the spontaneous blink generator: an animal model. The Journal of neuroscience: the official journal of the Society for Neuroscience 2011;31(31):11256–11267.

[31] Kandasamy R, Calsbeek JJ, Morgan MM. Home cage wheel running is an objective and clinically relevant method to assess inflammatory pain in male and female rats. J Neurosci Methods 2016.

[32] Kandasamy R, Lee AT, Morgan MM. Depression of home cage wheel running is an objective measure of spontaneous morphine withdrawal in rats with and without persistent pain. Pharmacology, biochemistry, and behavior 2017;156:10–15.

[33] Ko JA, Mizuno Y, Ohki C, Chikama T, Sonoda KH, Kiuchi Y. Neuropeptides released from trigeminal neurons promote the stratification of human corneal epithelial cells. Investigative ophthalmology & visual science 2014;55(1):125–133.

[34] Koltzenburg M, Wall PD, McMahon SB. Does the right side know what the left is doing? Trends Neurosci 1999;22(3):122–127.

[35] LaVail JH, Johnson WE, Spencer LC. Immunohistochemical identification of trigeminal ganglion neurons that innervate the mouse cornea: relevance to intercellular spread of herpes simplex virus. JComp Neurol 1993;327(1):133–140.

[36] Levine JD, Fields HL, Basbaum AI. Peptides and the primary afferent nociceptor. Journal of Neuroscience 1993;13:2273–2286.

[37] Levitt AE, Galor A, Weiss JS, Felix ER, Martin ER, Patin DJ, Sarantopoulos KD, Levitt RC. Chronic dry eye symptoms after LASIK: parallels and lessons to be learned from other persistent post-operative pain disorders. Molecular pain 2015;11:21.

[38] Ma F, Zhang L, Oz HS, Mashni M, Westlund KN. Dysregulated TNFalpha promotes cytokine proteome profile increases and bilateral orofacial hypersensitivity. Neuroscience 2015;300:493–507.

[39] Marfurt CF, Cox J, Deek S, Dvorscak L. Anatomy of the human corneal innervation. ExpEye Res 2010;90(4):478–492.

[40] Matzner O, Devor M. Hyperexcitability at sites of nerve injury depends on voltage-sensitive Na+ channels. J Neurophysiol 1994;72(1):349–359.

[41] Meng ID, Kurose M. The role of corneal afferent neurons in regulating tears under normal and dry eye conditions. Experimental eye research 2013;117:79–87.

[42] Muller LJ, Marfurt CF, Kruse F, Tervo TM. Corneal nerves: structure, contents and function. ExpEye Res 2003;76(5):521–542.

[43] Nakamura A, Hayakawa T, Kuwahara S, Maeda S, Tanaka K, Seki M, Mimura O. Morphological and immunohistochemical characterization of the trigeminal ganglion neurons innervating the cornea and upper eyelid of the rat. JChemNeuroanat 2007;34(3–4):95–101.

[44] Nosch DS, Pult H, Albon J, Purslow C, Murphy PJ. Relationship between Corneal Sensation, Blinking, and Tear Film Quality. Optometry and vision science: official publication of the American Academy of Optometry 2016;93(5):471–481.

[45] Parra A, Madrid R, Echevarria D, del Olmo S, Morenilla-Palao C, Acosta MC, Gallar J, Dhaka A, Viana F, Belmonte C. Ocular surface wetness is regulated by TRPM8-dependent cold thermoreceptors of the cornea. Nature medicine 2010;16(12):1396–1399.

[46] Peier AM, Moqrich A, Hergarden AC, Reeve AJ, Andersson DA, Story GM, Earley TJ, Dragoni I, McIntyre P, Bevan S, Patapoutian A. A TRP channel that senses cold stimuli and menthol. Cell 2002;108(5):705–715.

[47] Perini I, Tavakoli M, Marshall A, Minde J, Morrison I. Rare human nerve growth factor-beta mutation reveals relationship between C-afferent density and acute pain evaluation. J Neurophysiol 2016;116(2):425–430.

[48] Quallo T, Vastani N, Horridge E, Gentry C, Parra A, Moss S, Viana F, Belmonte C, Andersson DA, Bevan S. TRPM8 is a neuronal osmosensor that regulates eye blinking in mice. Nature communications 2015;6:7150.

[49] Ruohonen S, Jagodi M, Khademi M, Taskinen HS, Ojala P, Olsson T, Roytta M. Contralateral non-operated nerve to transected rat sciatic nerve shows increased expression of IL-1beta, TGF-beta1, TNF-alpha, and IL-10. J Neuroimmunol 2002;132(1–2):11–17.

[50] Simone DA, Nolano M, Johnson T, Wendelschafer-Crabb G, Kennedy WR. Intradermal injection of capsaicin in humans produces degeneration and subsequent reinnervation of epidermal nerve fibers: correlation with sensory function. The Journal of neuroscience: the official journal of the Society for Neuroscience 1998;18(21):8947–8959.

[51] Tran MT, Ritchie MH, Lausch RN, Oakes JE. Calcitonin gene-related peptide induces IL-8 synthesis in human corneal epithelial cells. Journal of immunology 2000;164(8):4307–4312.

[52] Tsujino H, Kondo E, Fukuoka T, Dai Y, Tokunaga A, Miki K, Yonenobu K, Ochi T, Noguchi K. Activating transcription factor 3 (ATF3) induction by axotomy in sensory and motoneurons: A novel neuronal marker of nerve injury. Mol Cell Neurosci 2000;15(2):170–182.

[53] Vashisht S, Singh S. Evaluation of Phenol Red Thread test versus Schirmer test in dry eyes: A comparative study. Int J Appl Basic Med Res 2011;1(1):40–42.

[54] Wang EF, Misra SL, Patel DV. In Vivo Confocal Microscopy of the Human Cornea in the Assessment of Peripheral Neuropathy and Systemic Diseases. BioMed research international 2015;2015:951081.

[55] Yagev R, Levy J, Shorer Z, Lifshitz T. Congenital insensitivity to pain with anhidrosis: ocular and systemic manifestations. Am J Ophthalmol 1999;127(3):322–326.

[56] Yamaguchi T, Hamrah P, Shimazaki J. Bilateral Alterations in Corneal Nerves, Dendritic Cells, and Tear Cytokine Levels in Ocular Surface Disease. Cornea 2016;35 Suppl 1:S65–S70.

[57] Yamaguchi T, Turhan A, Harris DL, Hu K, Pruss H, von Andrian U, Hamrah P. Bilateral nerve alterations in a unilateral experimental neurotrophic keratopathy model: a lateral conjunctival approach for trigeminal axotomy. PloS one 2013;8(8):e70908.

